# Generation of cell-free matrices that support human NK cell migration and differentiation

**DOI:** 10.1101/2020.01.26.920116

**Authors:** Barclay J. Lee, Emily M. Mace

## Abstract

Human natural killer cells are effectors of the innate immune system that originate from hematopoietic precursors in the bone marrow. While feeder cell lines that support NK cell development from hematopoietic precursors are often used to generate mature NK cells from lymphoid precursors in vitro, the nature of contributing factors of these stromal cells to the generation of functionally mature NK cells has been poorly described. Previous studies have shown that developing NK cells adhere to, and migrate on, developmentally supportive stroma. Here, we describe the generation of cell-derived matrices (CDMs) from a commonly used murine fetal liver stromal cell line. These CDMs are derived directly from the same EL08.1D2 stromal cell line known to support NK cell differentiation and contain ECM structural components fibronectin and collagen. We demonstrate that CDMs support NK cell adhesion and migration with similar properties as intact cells. Further, we show that CDMs support NK cell maturation from lymphoid precursors in vitro, albeit with reduced cell survival compared to intact cell-based differentiation. Together, these results describe a cell-free system that supports NK cell development and that can serve as a useful model for studying the nature of the biochemical interactions between NK cell developmental intermediates and developmentally supportive substrates.

## Introduction

Human natural killer (NK) cells originate from common lymphoid progenitors in the bone marrow, but likely undergo further maturation in peripheral tissues, the best described of which is secondary lymphoid tissue(1–4). While the contact-dependent signals that drive NK cell differentiation from precursors are not fully defined, this process can be recapitulated in vitro as a method to study NK cell development and ultimately define minimal requirements for the generation of mature, functional NK cells.

NK cell development in vitro frequently includes the use of stromal cell lines for co-culture with precursors in the presence of cytokines. Such stromal cell line co-culture methods lead to significant expansion of relatively mature NK cells that are functional for production of cytokines and lysis of susceptible target cells(5–8). The best characterized cell lines that support NK cell development are the OP9 or OP9-DL1 lines, derived from newborn mouse bone marrow (8), and EL08.1D2 murine fetal liver stromal cell lines (9). In addition, co-culture with bone marrow stroma, human splenic fibroblasts, mesenchymal stromal cells, and other stromal cell lines such as M2-10B4 and AFT024 can support NK cell differentiation from CD34^+^ precursors (10–14). Cell-free methods have also been designed and include the use of heparin-based Glycostem media, which functions to bind cytokines and help prevent their degradation while avoiding the use of xenogenic co-culture (15–17). While heparin-based methods can produce mature NK cells at the same frequency as stromal cell based methods, the use of stromal cells increases the expansion of developing NK cells and remains the most efficient means of generating the greatest number of mature cells (15). Notably, the effect of co-culture with supportive stromal cell lines is dependent upon direct cell-cell contact and the use of transwell barriers prevents NK cell development in these systems (5, 10).

Our previous studies sought to characterize the contact-dependent interactions between developing NK cells and these stromal cells, and we previously defined and characterized cell migration of NK cell developmental intermediates while undergoing differentiation and migration. We showed that NK cell migratory phenotype correlates with developmental stage and that NKDI derived in vitro from CD34^+^ stem cells exhibited progressively greater cell velocity and less frequent cell arrest as they matured (18, 19). With this knowledge, we hypothesized that cell-cell adhesions between NKDI and EL08.1D2 stroma drive cell migration while concurrently provide developmentally supportive signaling to promote an NK cell fate. We additionally observed that this cell migration was not disrupted following the loss of stromal cells from the co-culture and we hypothesized that this could be due to the production of extracellular matrix components from these cells that remained and supported the ongoing migration of NK cell developmental intermediates (19).

Here, we sought to generate and characterize cell-derived extracellular matrices from the EL08.1D2 stromal cell line. The generation of cell-free matrices has been previously described as a means of studying the mechanical properties of ECM, which is comprised of structural proteins such as collagen, fibronectin, and laminin, and serves as a scaffold for cells in vivo (20, 21). These studies have justified the use of cell-derived matrix as a model of cell activity within tissue as a preferable alternative to synthetic hydrogels such as Matrigel or collagen gels as they replicate the complexity of the ‘matrisome’ with greater fidelity (22). Here, we use ECM derived directly from EL08.1D2 as a model system in which only NK cell-ECM interactions are kept and determine the effect on cell adhesion, migration, and development. Using this model system, we partially elucidate requirements for NK cell differentiation and expansion, and identify ECM production by EL08.1D2 stroma as a contributing component of their support of NK cell development.

## Methods

### CD34^+^ precursor and primary NK cell isolation

T and B cell lineage depletion was performed using RosetteSep Human Hematopoietic Progenitor Cell Enrichment Cocktail (Stemcell Technologies) and Ficoll-Paque density gradient centrifugation from routine red cell exchange apheresis performed at Texas Children’s Hospital or Columbia University New York Presbyterian Hospital. Following pre-incubation with RosetteSep, apheresis product was layered on Ficoll-Paque for density centrifugation at 900 g (2,000 r.p.m.) for 20 min (no brake). Cells were harvested from the interface and washed with PBS by centrifugation at 1,500 r.p.m. for 5 min then resuspended in fetal calf serum for cell sorting. T- and B-cell depleted cultures were incubated with antibodies for CD34 (clone 561, PE conjugate, BioLegend, 1:100) prior to sorting. FACS sorting was performed using a BD Aria II cytometer with an 85 µm nozzle at 45 p.s.i. Purity after sorting was >80%. Primary NK cells for short-term imaging were similarly enriched with NK cell RosetteSep.

For FACS analysis of CD34^+^ intermediates, a 6-colour flow cytometry panel was performed on a Bio-Rad ZE5 Cell Analyzer using antibodies for CD56 (Clone HCD56, Brilliant Violet 605, BioLegend, 1:200), CD3 (Clone SK7, Brilliant Violet 711, BioLegend, 1:100), CD16 (Clone 3G8, PE-CF594 conjugate, BD, 1:300), CD94 (Clone DX22, APC conjugate, BioLegend, 1:100), CD117 (Clone 104D2, PE/Cy7 conjugate, BioLegend, 1:10), and a Zombie Aqua Viability Dye (1:100, BioLegend). Flow cytometry data analysis was performed with FlowJo X (TreeStar Inc.).

### Cell culture

EL08.1D2 cells stromal cells (a gift from Dr. J. Miller, University of Minnesota) were maintained on gelatinized culture flasks at 32° C in 40.5% α-MEM (Life Technologies), 50% Myelocult M5300 (STEMCELL Technologies), 7.5% heat-inactivated fetal calf serum (Atlanta Biologicals) with β-mercaptoethanol (10^−5^ M), Glutamax (Life Technologies, 2 mM), penicillin/streptomycin (Life Technologies, 100 U ml^−1^), and hydrocortisone (Sigma, 10^−6^ M). Culture media was supplemented with 20% conditioned supernatant from previous EL08.1D2 cultures.

NK92 cells (ATCC) were maintained in 90% Myelocult H5100 (Stemcell Technologies) with IL-2 (200 U ml^−1^). Cell lines were authenticated by flow cytometry and confirmed monthly to be mycoplasma free.

### CDM preparation

To prepare plates for CDM production, 96-well cell culture plates were treated with 0.1% gelatin in ultrapure water. 50 µL of 0.1% gelatin was deposited on 96-well plates and incubated at 37°C for 1 hour. Following incubation, wells were washed with sterile PBS. 50 uL of 1% (vol/vol) glutaraldehyde (Sigma-Aldrich) was then added for a 30 minute incubation at room temperature. After, wells were washed with PBS and treated with 1 M glycine (Sigma) for 2 min at room temperature. Wells were washed with PBS and stored overnight at 4°C in EL08.1D2 cell culture media supplemented with 1% P/S (Life Technologies).

EL08.1D2 cells were trypsinized and added to the gelatinized 96-well plates at a density of 10^4^–10^5^ cells/cm^2^ in a 150 µL volume. Cells were incubated at 32°C until formation of a confluent monolayer. Once a monolayer formed, culture media was replaced with a solution of culture media supplemented with ascorbic acid (50 µg ml^−1^; Sigma). Media was replaced every second day with freshly made ascorbic acid supplemented media for 7-10 days.

After this treatment, cells were extracted by aspirating off media and washing the cells with PBS. Stromal cells were denuded by adding 50 µL of extraction solution consisting of 2% (vol/vol) NH4OH (Sigma), 0.5% (vol/vol) Triton X-100 (Fisher) in PBS, which was first pre-warmed to 37°C. After a 3 minute incubation at room temperature, extraction buffer was pipetted off and wells were washed twice with PBS. Next, 50 µL of DNase I (10 µg ml^−1^; Sigma-Aldrich) was deposited and incubated at 37°C for 30 minutes. Then the DNase solution was removed, followed by washing twice with PBS. The resulting CDMs were stored at 4°C in PBS supplemented with 1% (vol/vol) penicillin-streptavidin.

### In vitro differentiation from CD34^+^ hematopoietic precursors

For in vitro CD34^+^ differentiation on EL08.1D2, 96-well plates were treated with 0.1% gelatin in ultrapure water to promote cell adherence. 50 µL of 0.1% gelatin was deposited on 96-well plates and incubated at room temperature for 30 minutes. Following incubation, wells were washed with PBS and left for an additional 60 minutes in a sterile culture hood to dry. Gelatinized 96-well plates were pre-coated with a confluent layer of EL08.1D2 cells at a density of 5-10 × 10^3^ cells per well and then mitotically inactivated by irradiation at 300 rad. For in vitro CD34^+^ differentiation on CDM, 96-well plates containing cell-derived matrix were prepared as described above. For in vitro differentiation on low adhesion plates, low adhesion plates (Corning) were used without additional preparation.

Purified CD34^+^ hematopoietic precursors were cultured at a density of 2-20 × 10^3^ cells per well on EL08.1D2 or CDM plates in Ham F12 media plus DMEM (1:2) with 20% human AB-serum, ethanolamine (50 µM), ascorbic acid (20 mg ml^−1^), sodium selenite (5 µg ml^−1^), β-mercaptoethanol (24 µM) and penicillin/streptomycin (100 U ml^−1^) in the presence of IL-15 (5 ng ml^−1^), IL-3 (5 ng ml^−1^), IL-7 (20 ng ml^−1^), Stem Cell Factor (20 ng ml^−1^), and Flt3L (10 ng ml^−1^) (all cytokines from Peprotech). Half-media changes were performed every 7 days, excluding IL-3 after the first week.

### Western Blotting

EL08.1D2 cells were lysed in RIPA buffer (Thermo Fisher) with 1% vol/vol HALT inhibitor (Thermo Fisher) at a concentration of 10^7^ cells ml^−1^. Samples were mixed with LDS buffer and DTT and separated on a 4-12% Bis-Tris gel. Proteins were transferred to a nitrocellulose membrane then blocked with skim milk. For CDM western blots, matrix was scraped directly into LDS/DTT solution and pre-heated at 95°C for 5 minutes. Fibronectin was detected with a polyclonal rabbit anti-fibronectin antibody (1:1000; Abcam) and actin with mouse anti-actin (1:5,000; Sigma) followed by either goat anti-mouse 680 or goat anti-rabbit 800 (1:10,000; LiCor). Proteins were detected on a LiCor Odyssey.

### Acquisition of microscopy images

For tracking of NK developmental intermediates, cells were seeded at 2 × 10^3^ cells per well on a 96-well ImageLock plate with confluent irradiated EL08.1D2 cells, then imaged at 2 minute intervals on the IncuCyte ZOOM Live-Cell Analysis System (Essen Bioscience) at 37° C in the phase-contrast mode (10X objective). Images were acquired continuously for a 21-day period and data were exported as TIFF image series for further analyses.

For microscopy, EL08.1D2 cells or cell-derived matrices were prepared on circular microscope coverslips in 24-well cell culture plates. Plates were blocked with heat denatured 2% BSA/PBS for 1 hour at 37 °C. Primary NK cells from lineage-depleted apheresis or NK92 cells were incubated on cells or cell-free matrix for 30 minutes in the presence of directly conjugated antibodies to surface antigens CD56 (1:50, clone HCD56, Biolegend) and CD29 (1:50, clone TS2/16, BioLegend). Samples were fixed with 4% PFA/PBS for 20 minutes at room temperature, then washed three times with PBS and treated with 0.2% Triton X-100/PBS for 10 minutes. Samples were stained with polyclonal rabbit anti-collagen I antibody (1:100, Novus Biologicals), followed by phalloidin (Thermo Fisher). Samples were washed and coverslips were removed and mounted onto microscope slides in ProLong Gold (Life Technologies). Images were acquired on a Zeiss Axio Observer Z1 microscope with a Yokigawa W1 spinning disk and a 63X 1.40 NA objective. Illumination was by solid state laser and detection by Prime 95B sCMOS camera. Data were acquired in 3i Slidebook software and exported to Fiji (23) for further analysis.

### Image analysis and cell tracking

For analysis of collagen confluence, microscopy images showing collagen distribution were imported into Fiji, and manually thresholded to remove background. The confluence was then recorded based on the ‘Area Fraction’ measurement. For actin footprint measurement, a freehand selection was drawn around cells and the area was then measured.

Cell tracking from Incucyte image series was performed using the TrackMate plugin in Fiji by manual tracking (24). Track statistics were exported from Fiji and the data was imported into MATLAB, which was also used to generate rose plots. Mean speed, straightness, and arrest coefficient were calculated using custom scripts in MATLAB. Mean speed was defined as the per-track average of all instantaneous speeds calculated at each frame. Accumulated distance was measured as the cumulative length of each track measured within a single imaging series. The straightness parameter was calculated by dividing the displacement by the total path length for each track. Arrest coefficient was defined as the percentage of time that the cell stays in arrest based on a threshold on instantaneous speed of 2 µm/min, or approximately one cell body length per image interval.

### Statistics

Statistical analysis was calculated using Prism 8.0 (GraphPad). Ordinary one-way ANOVA was used to compare track statistics. Mann-Whitney U test was used to compare actin footprint data. For all tests P<0.05 was considered significant.

## Results

### Cell-derived matrices from EL08.1D2 contain common ECM components and support NK cell adhesion

Given our previous observation that cells migrated on previously EL08.1D2 confluent surfaces, we sought to define whether EL08.1D2 secrete extracellular components. Cell-derived matrices (CDMs) were generated from EL08.1D2 stroma using modified protocols from previously described methods (20, 21, 25). Briefly, cell culture plates were coated with gelatin to promote cell adhesion. As gelatin is soluble in aqueous solution, plates were then treated with glutaraldehyde to crosslink the gelatin and enhance its stability in solution (26). EL08.1D2 cells were added to the plates and allowed to adhere overnight. After a confluent monolayer was formed, ascorbic acid was cycled into the culture media for a period of 7 days in order to promote CDM formation. Following this, the cellular and nuclear debris were removed, leaving a completed cell-derived matrix that could then be used for further experiments (summarized in Fig. 1).

**Fig. 1.**
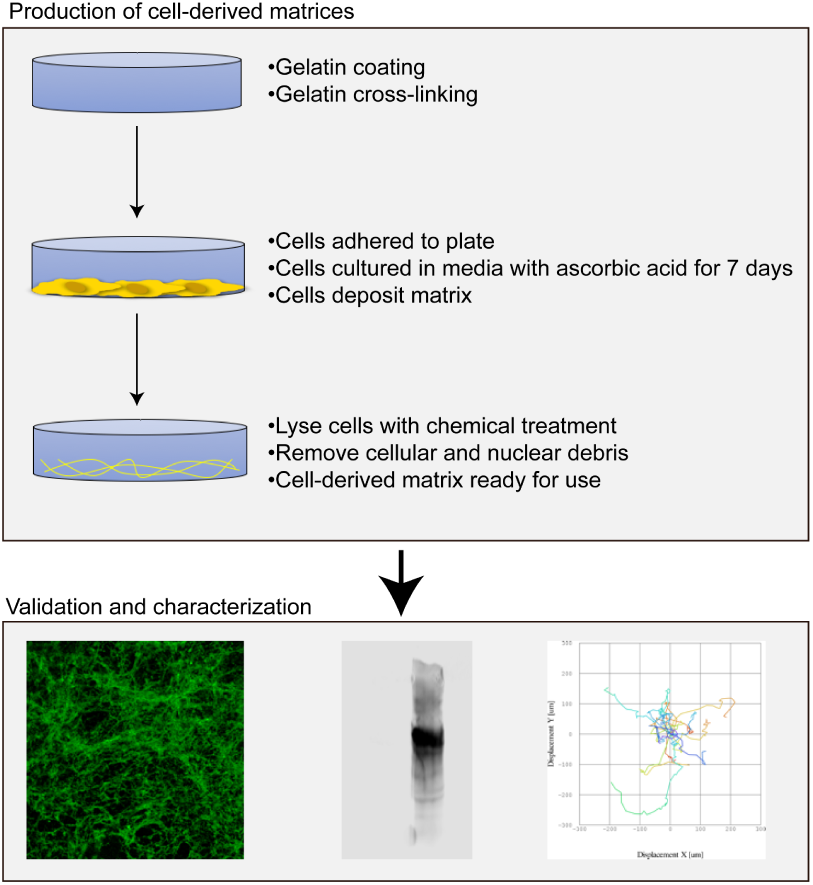
Workflow for generation of cell-free matrices from EL08.1D2 stroma. An overview of the protocol for the generation of CDM. Cell culture plates are coated with gelatin and seeded with EL08.1D2 stromal cells. Cells are allowed to adhere overnight and grown until confluent. Once confluent, ascorbic acid is cycled into the media for 7 days to promote matrix formation. During matrix extraction, cells are lysed with a chemical treatment and nuclear debris is removed with DNAse. CDMs can then be validated by microscopy and western blotting, or used for functional studies such as NK cell migration.

To determine whether this culture method was inducing the secretion of ECM components from EL08.1D2 cells, CDMs were stained with collagen I antibody and imaged by confocal microscopy. Analysis of microscopy images revealed that cell-derived matrices contained a collagen network that was significantly more confluent than that of intact EL08.1D2 stromal cells, with a mean confluence of 13.76±9.99% versus 0.29±0.50% (Fig. 2A, B). As expected, control microscopy slides without EL08.1D2 stroma did not display a collagen network, even after treatment for CDM production, with mean confluences of 0.009±0.016% and 0.467±0.552% for cell-free controls with and without CDM treatment, respectively. To further confirm these results, we performed Western blots on solubilized CDM and EL08.1D2 cell lysates for fibronectin and actin content. Indeed, fibronectin content was detected on solubilized CDM but not on EL08.1D2 lysates (Fig. 2C). These experiments, which additionally detected actin in lysates from intact EL08.1D2 cells but not CDM lysate, demonstrated that cell-derived matrices were being produced from EL08.1D2 stroma.

**Fig. 2.**
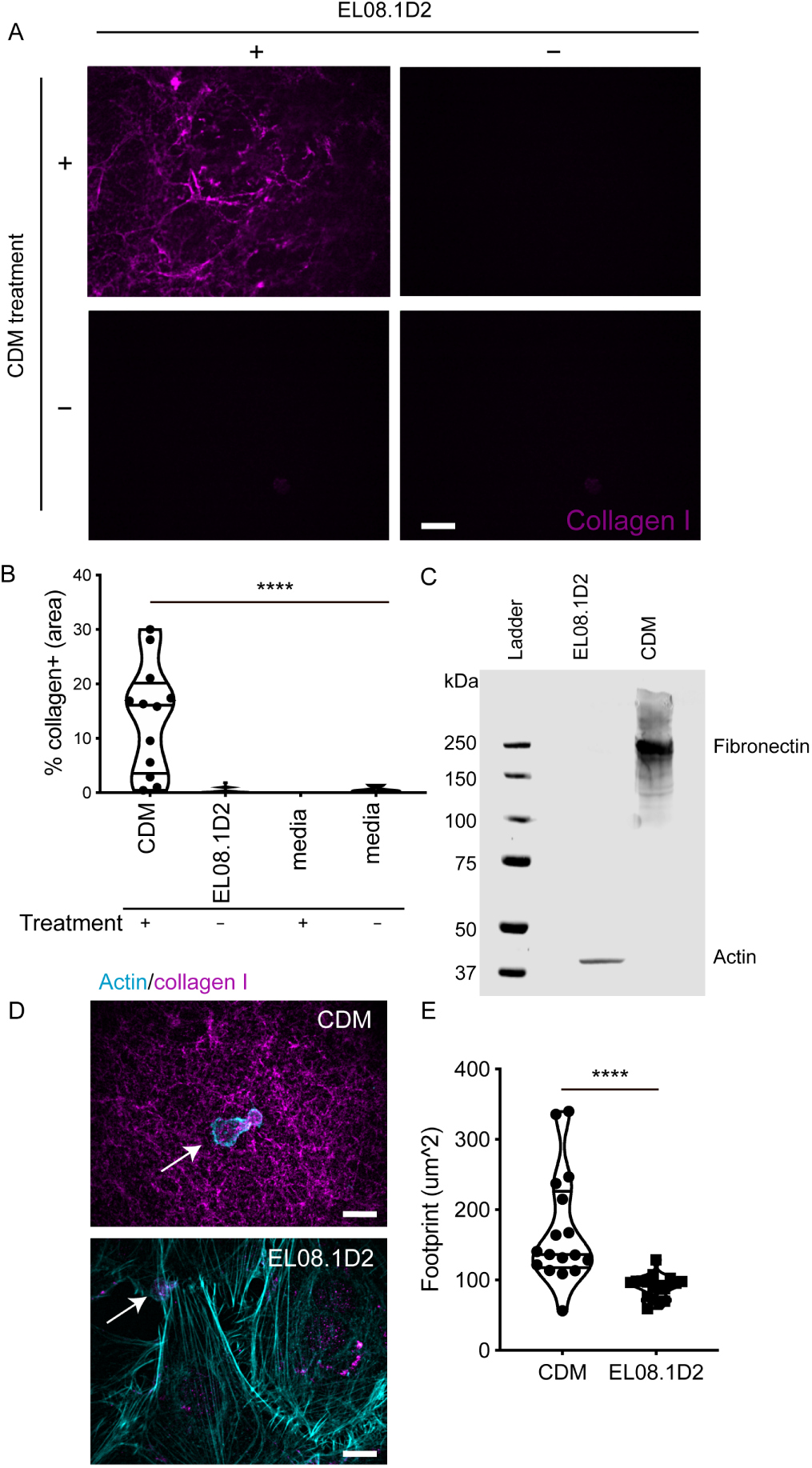
Cell-derived matrices contain ECM components and support NK cell adhesion. CDMs were generated as described in the text and compared with confluent intact EL08.1D2 cells or untreated control conditions. A) Fluorescence microscopy images of conditions used to validate CDM production. Only slides containing EL08.1D2 that underwent treatment for CDM production (top left) produced a collagen network. Slides with EL08.1D2 that were untreated (bottom left), slides without EL08.1D2 that underwent treatment for CDM production (top right), and slides without EL08.1D2 that were untreated (bottom right) did not generate detectable collagen. B) The area of the collagen network was measured for each condition and expressed as a frequency of the total field of view. Error bars represent s.d. Means with significant differences were determined by one-way ANOVA with Tukey’s multiple comparison test (****p< 0.0001). n=12-15 fields of view for CDM and EL08.1D2 conditions, 3-5 fields of view for negative controls. C) CDM were solubilized in lysis buffer or cell lysates were prepared from intact EL08.1D2 stromal cells. Proteins were separated by SDS-PAGE and fibronectin or actin were detected by Western blotting. D) Confocal microscopy images showing NK92 adhered to CDM or intact EL08.1D2 cells following immunostaining for collagen (magenta) or actin (cyan). Arrows denote NK92 cells, images shown as maximum projection of 29 Z slices. Scale bar=10 µm. E) Quantification of cell footprint in each condition. Error bars indicate s.d. Means with significant differences were determined by Mann-Whitney U test (****p< 0.0001). n=17 measurements per condition.

To determine if CDMs supported NK cell adhesion, we imaged the human NK cell line NK92 cells incubated on CDM or EL08.1D2 stroma. NK92 were seeded on each substrate, allowed to adhere for 30 minutes, and then fixed and stained with antibodies against actin and collagen (Fig. 2D). In addition to the significant detection of collagen described for CDM above, EL08.1D2 stroma also displayed a minor but detectable level of collagen expression. Confocal microscopy revealed that NK92 adhered to both CDM and EL08.1D2 stroma, although NK92 on CDMs seemed to display greater spreading, including long, thin filopodia-like extensions. Quantitative image analysis confirmed that NK92 cells exhibited a significantly larger actin footprint on CDMs compared to EL08.1D2 stroma, with a mean footprint of 165.0±60.4 µm2 on CDMs compared to 90.2±16.9 µm^2^ on stroma (Fig. 2E).

Together, these data demonstrate that EL08.1D2 secrete ECM proteins that form collagen and fibronectin-containing matrices that supports human NK cell adhesion and spreading.

### CDMs support adhesion and migration of primary NK cells

Having confirmed that CDMs supported NK92 adhesion, we repeated these experiments with freshly isolated ex vivo NK cells to determine if primary cells also adhered to CDM. Ex vivo NK cells were isolated from human peripheral blood, incubated on CDM or EL08.1D2 stroma, and fixed and stained with antibodies against actin, integrin β1 (CD29), and CD56. Confocal microscopy of cells again showed the presence of filopodia-like extensions characterized by integrin β1 and CD56 enrichment, demonstrating that integrin β1 is important for adhesion to both stroma and ECM (Fig. 3A). Z-stack information revealed that CDMs were somewhat 3-dimensional with a depth comparable to that of the EL08.1D2 monolayer as previously reported (27), and NK cells were capable of invading into the CDM. Once again, primary NK cells had greater spreading on CDMs, with a mean actin footprint of 40.0±12.3 µm2^2^ on CDM compared to 36.2±11.8 µm^2^ on stroma (Fig. 3B).

**Fig. 3.**
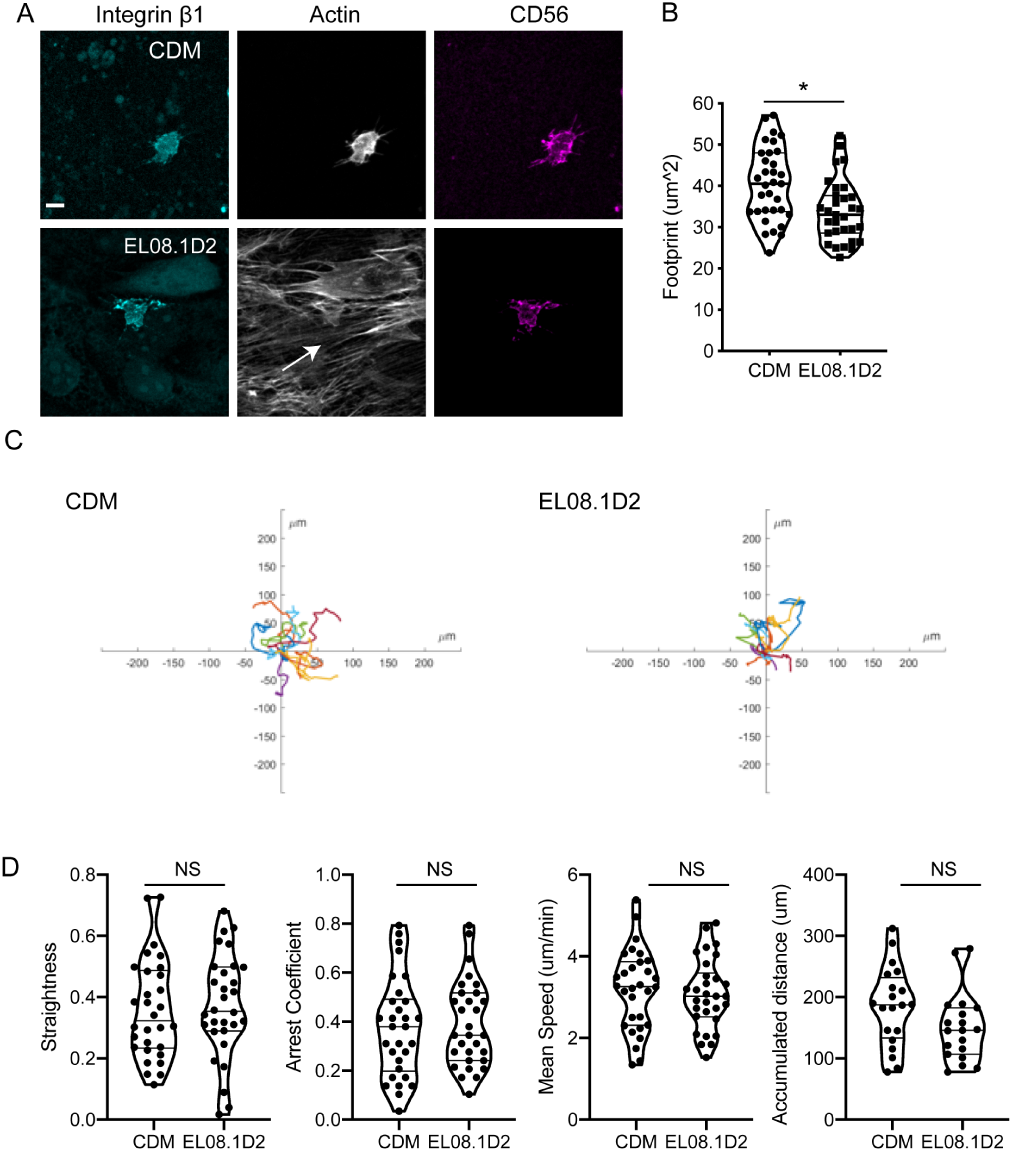
Primary NK cells adhere to and migrate on cell-free matrices. Freshly isolated primary NK cells were adhered to either EL08.1D2 or cell-derived matrix. A) cells were co-incubated for 30 minutes and then fixed and immunostained as indicated followed by confocal microscopy. Images show maximum intensity projection of z-stacks. Scale bar=10 µm. B) Quantification of cell area in each condition. Error bars indicate s.d. Means with significant differences were determined by Mann-Whitney U test (*p< 0.05). C) Primary NK cells were isolated from peripheral blood and seeded on substrates and imaged every 2 minutes. Cell migration was tracked following image acquisition. Rose plots of primary NK cell tracks in each condition are shown. D) Cells were tracked and migratory parameters were measured. n=19, 20 cells from 2 individual healthy donors. Means were analyzed by Mann-Whitney test.

As CDMs supported adhesion of both primary NK and NK92 cells, we hypothesized that cell migration of primary NK cells on CDM would be comparable to that of cells on stroma. Primary cells were seeded on either CDM, EL08.1D2, or low-adhesion plates coated with a hydrophilic, neutrally charged hydrogel to inhibit cell attachment. Immediately after cell addition, live-cell imaging was performed with image acquisition every 2 minutes. Cells were tracked from live cell imaging series (Fig. 3C). We found that freshly isolated primary NK cells had comparable migratory phenotypes on CDM and EL08.1D2 (Fig. 3D). In contrast, cells did not migrate or adhere on low adhesion plates and cell motions that were detected in imaging were predominantly drift of non-adherent cells (not shown). These results demonstrate that CDMs support integrin-dependent adhesion and migration of primary NK cells.

### CDMs can partially support NK cell development from hematopoietic precursors

Having established that CDMs support NK cell adhesion and migration, and given that EL08.1D2 stroma support NK cell development, we sought to assess NK development on CDMs. CD34^+^ hematopoietic precursors were isolated from peripheral blood and seeded on either CDM, stroma, or low adhesion plates. Flow cytometry analysis for developmental markers was performed weekly. At 21 days, NKDI showed similar maturation on EL08.1D2 and CDMs but failed to survive on low adhesion plates (Fig. 4A). Expression of CD56 and CD94 was consistently observed for the first two conditions, denoting that NK cell developmental intermediates had successfully reached stage 4 of NK development. However, NK cell receptor expression was slightly decreased in the CDM condition, with 49.1% of NKDIs positive for CD94 compared to 65.9% of NKDIs positive in the EL08.1D2 condition (Fig. 4B). Similarly, only 63% of NKDIs in the CDM condition were positive for CD56 expression, whereas 80.3% of NKDIs were positive in the EL08.1D2 condition.

**Fig. 4.**
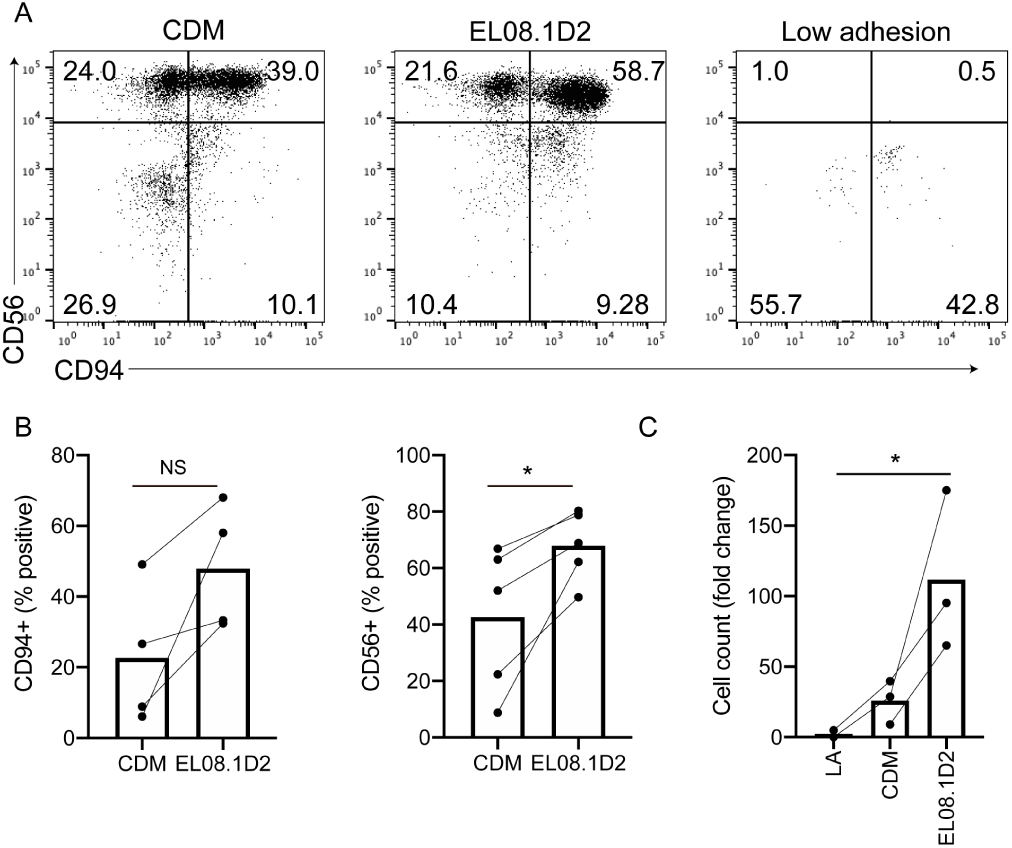
CDM partially support NK cell development. CD34^+^ HSCs were isolated from peripheral blood and seeded on CDM, EL08.1D2, or low adhesion plates. A) After 21 days cells were harvested and analyzed by flow cytometry to determine cell maturation. Cells were gated on FSC/SSC and a live-dead marker for live lymphocytes. Representative flow cytometry plots are shown from 2 independent repeats with different donors. B) Percentage of cells positive for CD94 or CD56 following culture of CD34+ precursors on paired CDM and EL08.1D2 conditions. n=4 (CD94), 5 (CD56). *p<0.05 by two-tailed paired t-test. C) Cell expansion shown as fold change from number of CD34^+^ precursors seeded at day 0. n=2 (low adhesion), 3 (CDM and EL08.1D2) independent replicates from different healthy donors. *p<0.05 by ordinary one-way ANOVA.

Previous studies have identified the promotion of cell survival and proliferation by EL08.1D2 as a key contributing factor to their efficiency at generating mature NK cells (15). In keeping with these data, cell counts showed that NK survival and proliferation was decreased in the CDM condition (Fig. 4C). Together, these results suggest a requirement for adhesion/migration for NK cell development; HSCs on low adhesion plates failed to develop into mature NK cells in contrast to those plated on CDM or EL08.1D2, which proliferated and matured. However, both cell maturation and survival of NKDI on CDM was inferior to that of the EL08.1D2 plates, suggesting that adhesion is necessary but not sufficient for NK development.

### NK cell developmental intermediates undergo migration on CDM and EL08.1D2

Given that NKDI were capable of maturing on cell-derived matrices, we hypothesized that they would have similar migrational behavior on CDM as has been previously described for cells undergoing maturation on EL08.1D2 (18, 19). CD34^+^ hematopoietic precursors were isolated from peripheral blood and seeded on either CDM or EL08.1D2. In conjunction with the weekly flow cytometry for developmental markers, cells were also imaged every 2 minutes and cell migration was tracked at weekly time points (Fig. 5A). We found that the properties of tracks did not significantly differ between the two conditions at each weekly time point measured (Fig. 5B-D). For example, the mean speed of tracks was 2.14 ± 0.79 µm/min on EL08.1D2 after 0 days, and 1.93 ± 0.98 µm/min on CDMs (Fig. 5C). After 21 days, this increased to 3.626 ± 0.72 µm/min on EL08.1D2 and 2.936 ±0.79 µm/min on CDMs; this was consistent with previously reported migration speeds of mature NK cells in similar conditions (18, 27).

**Fig. 5.**
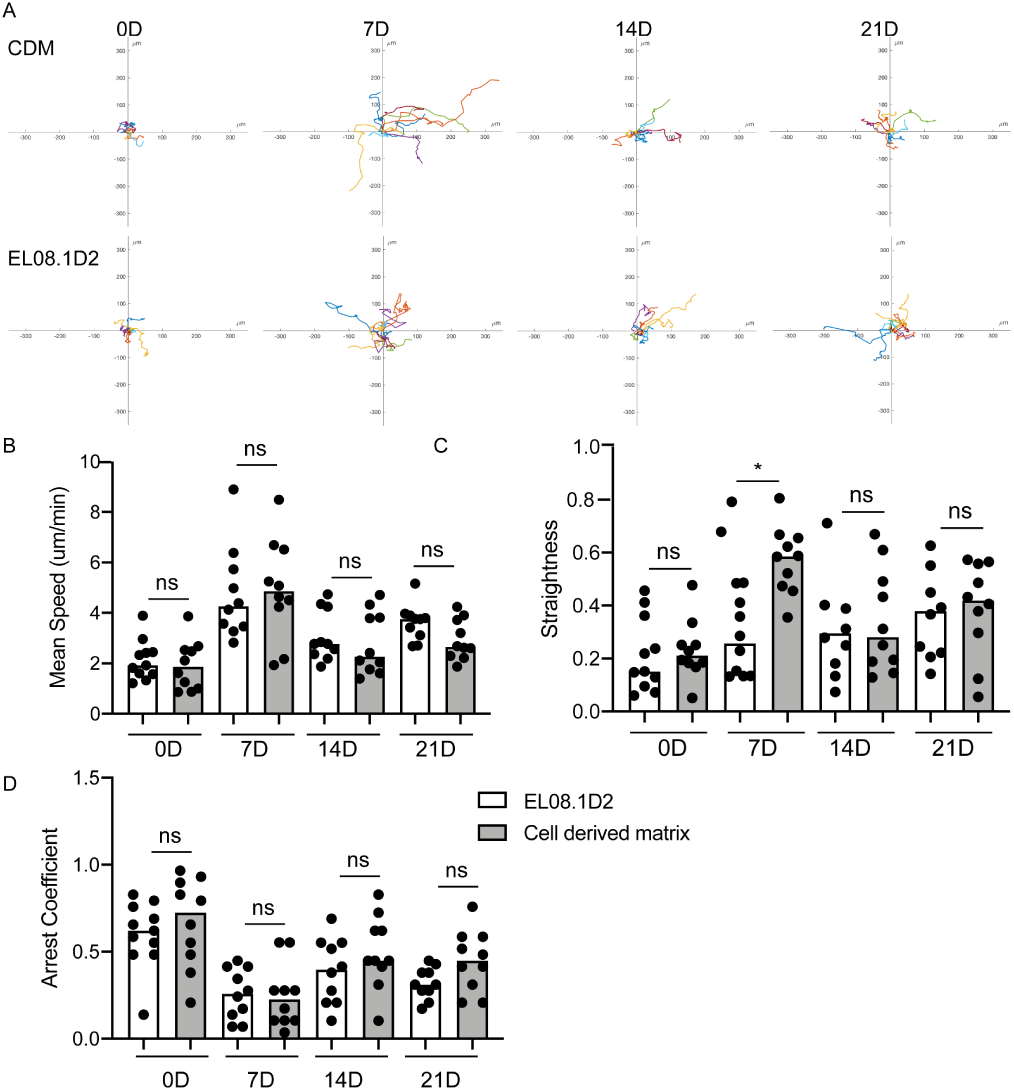
NKDIs have comparable migration on CDM and EL08.1D2. CD34^+^ HSCs were isolated from peripheral blood and seeded on CDM or EL08.1D2 stroma. Cell migration was imaged every 2 minutes and cells tracks were determined at each week. A) Representative tracks showing HSC migration on CDM and EL08.1D2 stroma. Scale bar=50 µm. B) Rose plots of cell tracks on CDM and EL08.1D2 at each week. C) At each week, the mean speeds (left), straightnesses (middle), and arrest coefficient (right) of cell tracks was determined. Error bars indicate s.d. Means with significant differences were determined by ordinary one-way ANOVA with Tukey’s multiple comparison test (* p< 0.05).

Similarly, after 0 days, mean straightness was 0.21 ± 0.14 on EL08.1D2 and 0.23 ± 0.11 on CDM, and mean arrest coefficient was 0.60 ± 0.19 on EL08.1D2 and 0.67 ± 0.26 on CDM (Fig. 5C, D). The only parameter in which we detected significant differences was straightness after 7 days, which was significantly higher on CDMs, with a mean straightness of 0.57 ± 0.13 compared to 0.28 ± 0.15 on EL08.1D2. This difference was relatively transient, however, as cells at 14 and 21 days had similar straightness properties. Taken together with our primary NK cell and cell line data, these data support the observation that CDM can support the migration of both mature and immature NK cells in a similar capacity as intact cells.

## Discussion

The use of stromal-based cell co-culture methods as a means of generating mature NK cells has been well described and has the potential to improve the immunotherapy approaches and provide insights into requirements for human NK cell development. However, the biochemical, physical and mechanical contributions of stromal cells to this process has not been fully described. Here, we describe the generation of fibronectin- and collagen-containing cell-derived matrices from EL08.1D2 stromal cells that can act as a substrate for NK cell adhesion and migration. We further show that these matrices can partially support in vitro NK cell development but with impaired expansion of NK cell developmental intermediates. These data extend our understanding of the nature of the contribution of stromal cells to lymphocyte development and support previous observations that stromal cells could be contributing ECM components to co-cultures with NK cell developmental intermediates (19).

Cell-free matrices have been well-described as a powerful tool for the studies of cell migration, tumor properties and tissue regeneration (20, 21, 28). They provide a useful model that mimics both the complex composition and substrate stiffness (29) of the in vivo cellular environment, while providing a substrate that is highly amenable to imaging by light microscopy and studies of cell motility and invasion. CDMs derived from various cell types have unique ECM components that reflect their unique matrisomes, however key ECM proteins including collagen, laminins and fibronectin are conserved. While here we limited our characterization of CDMs to the presence of collagen and fibronectin, future studies will focus on higher-resolution analyses of both the matrix components and secreted factors from supportive and non-supportive cell lines.

Our detection of collagen and fibronectin in EL08.1D2-derived CDM, in conjunction with the adhesion and migration of primary NK cells and NK cell developmental intermediates, suggests that integrins are mediating interactions between cells and CDMs. Integrins are heterodimeric receptors that bind a number ECM components, and the asparagine-glycine-aspartic acid (RGD) peptide is the basis of recognition of fibronectin by as many as 12 of the >20 known integrins. While the expression of integrins on hematopoietic cells is more restricted than that of adherent cells, human NK cells and developmental intermediates express integrin β1 and α4β1 (VLA-4) and α5β1 (VLA-5) mediate adhesion of freshly isolated human NK cells to fibronectin (30). This is consistent with a dependency of adherent cells on β1 integrin for adhesion and migration in 3D cell-free matrices (28). The increased cell spreading that we observed following short-term incubation of cells on CDMs relative to intact cells suggests that the properties that mediate this adhesion may differ from those of intact EL08.1D2 cells. This observation could reflect a relatively greater density of collagen and fibronectin on the CDMs when compared to intact cells, it could also be indicative of greater substrate stiffness induced by the glutaraldehyde cross-linking step (29). Regardless, it is notable that both primary and in vitro-derived NK cells migrated on CDM with no apparent differences in migratory properties from those cells migrating on intact EL08.1D2 cells. Specifically, we did not detect haptotaxis induced by matrix topology, as has been reported for cells migrating on fibroblast-derived ECM (20, 31, 32). This likely reflects differences in the composition or structure of the matrices between stromal cell-derived matrices and fibroblast-derived matrices, as well as different requirements for haptotaxis between immune cells and those that undergo non-amoeboid cell migration (33). Taken together with the adhesion data described above, however, the conserved patterns of migration seen on CDMs and intact cells suggests that the migration of NK cell developmental intermediates in an in vitro differentiation system can occur independently of cell-associated ligands, including ICAM and VCAM.

Whereas migration did not require the presence of intact cells, direct contact between NK cell developmental precursors and stroma greatly increased the expansion of precursors in our system as previously reported (15). The nature of the survival or proliferation signaling provided by stroma remains unknown, however previous studies have implicated conserved pathways such as Wnt and Notch, or secreted factors such as cytokines, in providing these signal (15). Additionally, bidirectional signaling between precursors and stroma may affect these interactions, as has been demonstrated for co-culture between EL08.1D2 and leukemic B cells (34). More explicit dissection of the effect of conditioned media or the use of previously co-cultured stroma would provide additional insight into the respective roles of soluble and contact-dependent signaling in these co-culture systems.

In summary, here we describe the generation of ECM components from the EL08.1D2 murine fetal liver cell line. Primary NK cells and NK cell developmental intermediates adhere to these cell-free matrices and undergo non-directed migration on these with similar properties to those on intact cells. Taken together, these data provide a greater understanding of the nature of developmental support provided by commonly used stromal feeder cells.

## ACKNOWLEDGEMENTS

This work was supported by NIH-NIAID R01AI137073 to EMM.

